# OmicsDB::Pathogens - A database for exploring functional networks of plant pathogens

**DOI:** 10.1101/2020.03.18.979971

**Authors:** Bjoern Oest Hansen, Stefan Olsson

**Affiliations:** State Key Laboratory for Ecological Pest Control of Fujian and Taiwan Crops, College of Plant Protection, Fujian Agriculture and Forestry University, Fuzhou 350002, China; OmicsDriven, Toelloese, Denmark; Max Planck Institute of Molecular Plant Physiology, Am Muehlenberg 1, 14476 Potsdam, Germany; Plant Immunity Center, Haixia Institute of Science and Technology, College of Life Science, Fujian Agriculture and Forestry University, Fuzhou 350002, China

## Abstract

Plant pathogens are a great threat to food security. To combat them we need an understanding of how they work. Integrating large-scale omics datasets such as genomes and transcriptomes has been shown to provide deeper insights into many aspects of molecular biology. For a better understanding of plant pathogens, we aim to construct a platform for accessing genomic and gene co-expression networks for a range of pathogens and reference species. Currently we have integrated genomic and transcriptomics data from 10 species (*Fusarium graminearum, Ustilago maydis, Blumeria graminis, Neurospora crassa, Schizosaccharomyces pombe, Saccharomyces cerevisiae, Escherichia coli, Arabidopsis thaliana, Mus musculus and Homo sapiens*).

Here we introduce OmicsDB::Pathogens (http://pathogens.omicsdb.org), a publicly available web portal with an underlying database containing genomic, and transcriptomic data and analysis tools. It allows non-bioinformaticians to browse genomic data and inspect and compare biological networks across species.

The information is modelled in a graph-based database, enabling flexibility for querying and future extensions. Tools such as BLAST and Cytoscape.js are available together with the option of performing GO enrichment analysis. The database also enables the user to browse information such as Orthologs, Protein domains and publications citing a given gene.

Herein we describe how to use this platform for generating hypotheses for the function of a gene.

**Availability and Implementation:** Currently, Omicsdb supports networks for 10 organisms and is freely available for public use at http://pathogens.omicsdb.org

## Introduction

Plant pathogens are a great threat to the ever-growing population and the following need for food. To combat them, an understanding of their mode of action and of how the pathogenic processes work is essential. To be able to obtain useful knowledge from genomes, it is necessary to be able to correctly assign functions to gene products. Manual annotation is not a feasible option to annotate the ever-growing number of available genomes. Scientists, therefore, must rely more and more on predictions.

In-silico methods have gained popularity for automated annotation of gene functions, though for it to be useful, it is important that the accuracy of the predictions, and number of genes that can be annotated is high.

The guilt by association hypothesis is based on the observation that genes that participate in similar biological processes or are involved in similar regulatory pathways tend to display similar expression profiles (Wolfe et al., 2005). An increasingly popular method to utilize this observation is using gene co-expression networks. Networks are a convenient way of representing biological data and allow for the utilization of graph theory. By analysing the structure of the networks, it is possible to identify communities within the network, where the genes are more closely related to each other than to the rest of the network and are enriched for specific biological processes.

The two main applications for co-expression network analysis are 1) To use a bait-gene from a known pathway or with a known function, as a query, and identify possible genes within the same pathway or with a similar function (Itkin et al., 2013) and 2) to suggest the biological process or pathways given gene is involved in, based on the functions of its neighborhood. With the increase in available biological networks, an increasingly important usage is the identification of common network patterns across species, this allows for more reliable transfer of knowledge from and across model species.

Inter-network comparisons serve multiple purposes, they can improve identification of functionally related orthologs across species (Hansen et al., 2014), and aid in the identification of conserved subgraphs within or across species (Stuart et al., 2003).

The amount of data a researcher working within molecular biology needs to handle has grown exponentially during later years. Not just does that put a heavy load on the IT infrastructure, but also on the researcher that might not be a bioinformatics specialist. Handling and processing omics data are often a requirement these days, to be able to perform analysis on this kind of data, researchers need to be familiar with scripting languages such as Python or R.

Curated databases have shown to be powerful tools for integration and analysis of data. This has led to an emergence of integrated web-based databases, changing the analysis of networks from being a task for specialist bioinformaticians, into a simple routine task for experimental molecular biologists, investigating specific genes or conditions. However, most of these databases focus on the major model species, such as Maize (Andorf et al., 2016), Budding yeast (Kim et al., 2014) or fission yeast (Vo et al., 2016). There exist very few options for researchers working on non-model species.

Plant pathogens have until now had little representation in these databases. Electronic resources for *Fusarium graminearum* eFG (Liu et al., 2013) a model species in plant pathology (Zhang et al., 2019), contains functional annotation as well as annotations of transcription factors and curated and predicted pathogenic genes for fusarium. PHI-base (Urban et al., 2017) is a great resource for curated Pathogen-Host Interactions but is manually curated and thus limited in size. PhytoPath (Pedro et al., 2016), is a resource for genomic information related to plant pathogens. Other databases with microbial data that cover mainly model species are also available (Kim et al., 2014, 2015; Oughtred et al., 2019)

To tackle this problem, we here present OmicsDB::Pathogens, an integrated database of networks from 8 species: 6 model species (*Escherichia coli, Saccharomyces cerevisiae, Saccharomyces pombe, Arabidopsis thaliana, Mus musculus, Homo sapiens*), and 4 plant pathogens (*Fusarium graminearum, Ustilago maydis, Blumeria graminis, Neurospora crassa and Magnaporthe oryzae*), as well as orthologs from 158 other fungal pathogens, to ease the knowledge transfer to and from these.

OmicsDB:: Pathogens is a database developed to assist experimental molecular biologists in accessing and analyzing omics data.

The database provides intuitive navigation, allowing the researchers to store, browse, analyse and compare their data, as well as handling metadata of both samples and workflows. Another important use of the database is that it allows comparisons between species that can allow the experimental researcher not just to infer possible functions of genes not previously studied experimentally in detail. It also makes it possible to rank genes of interest to study enabling more focussed research in sorting out gene function. We provide an extensive amount of already processed publicly available genomic and transcriptomic data.

## Results and discussion

OmicsDB is a database that includes high throughput experimental data and information for multiple species. Since we store data from biological networks, it was decided to go for the graph-based database Neo4J for storing most of the data. Neo4J has previously been evaluated by Have and Jensen for several graph processing problems related to bioinformatics and compared with PostgreSQL, they found Neo4J to be faster in many cases, though they also conclude that graph-based databases are not necessarily the best choice for all problems (Have and Jensen, 2013). Neo4J uses the property graph model, which means that nodes and edges can have key/value properties associated. Neo4J uses its own query language, Cypher for querying the graph, cypher allows queries to be formulated in terms of paths, which allows them to be concise and intuitive compared with equivalent SQL queries, which is often complicated by joins, and difficult to read. Due to the simplicity of traversing edges, and accessing data through the Cypher language, we decided to use Neo4J for all data gene expression data, that is stored in an SQLite database, and flat files that are stored in an S3 compatible object storage, with their paths and metadata stored in Neo4J linked to the relevant organism. A simplified overview of our Neo4J data model can be seen in Fig. 1 An overview of the infrastructure and data processing steps can be seen in Fig. 2. The processed expression data is stored in a SQL database, with common identifiers shared between the other systems.

**Fig 1.**
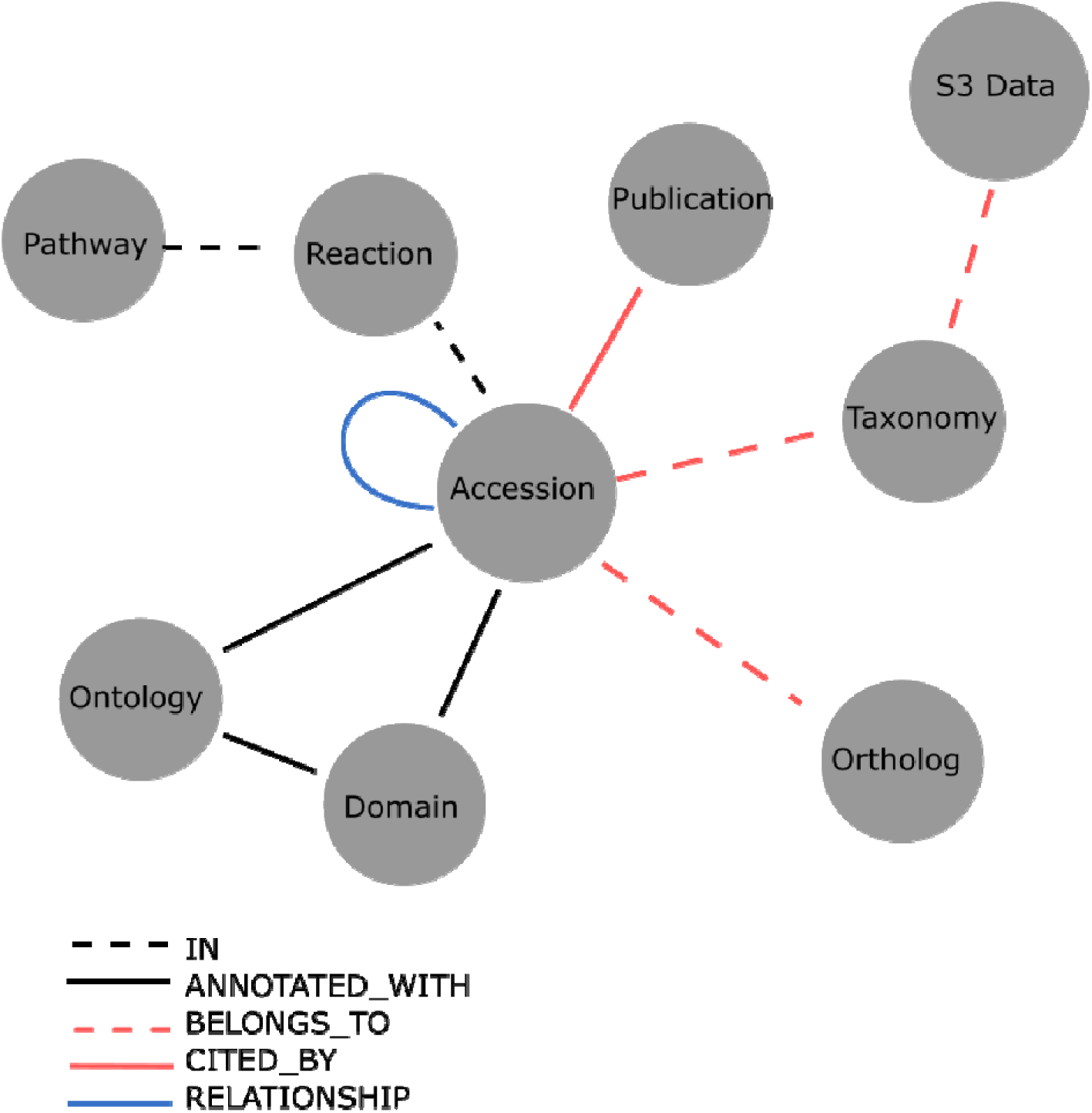
The part of our Neo4J data model. This shows how the different nodes are related. Properties are left out for simplicity. Gathering data across any node is easy, for example, if one wishes to know all publications related to a given Domain its possible to traverse the graph from Domain over Accession to Publication. We believe this gives a more intuitive way of accessing data than through a classical relational database.

**Figure 2.**
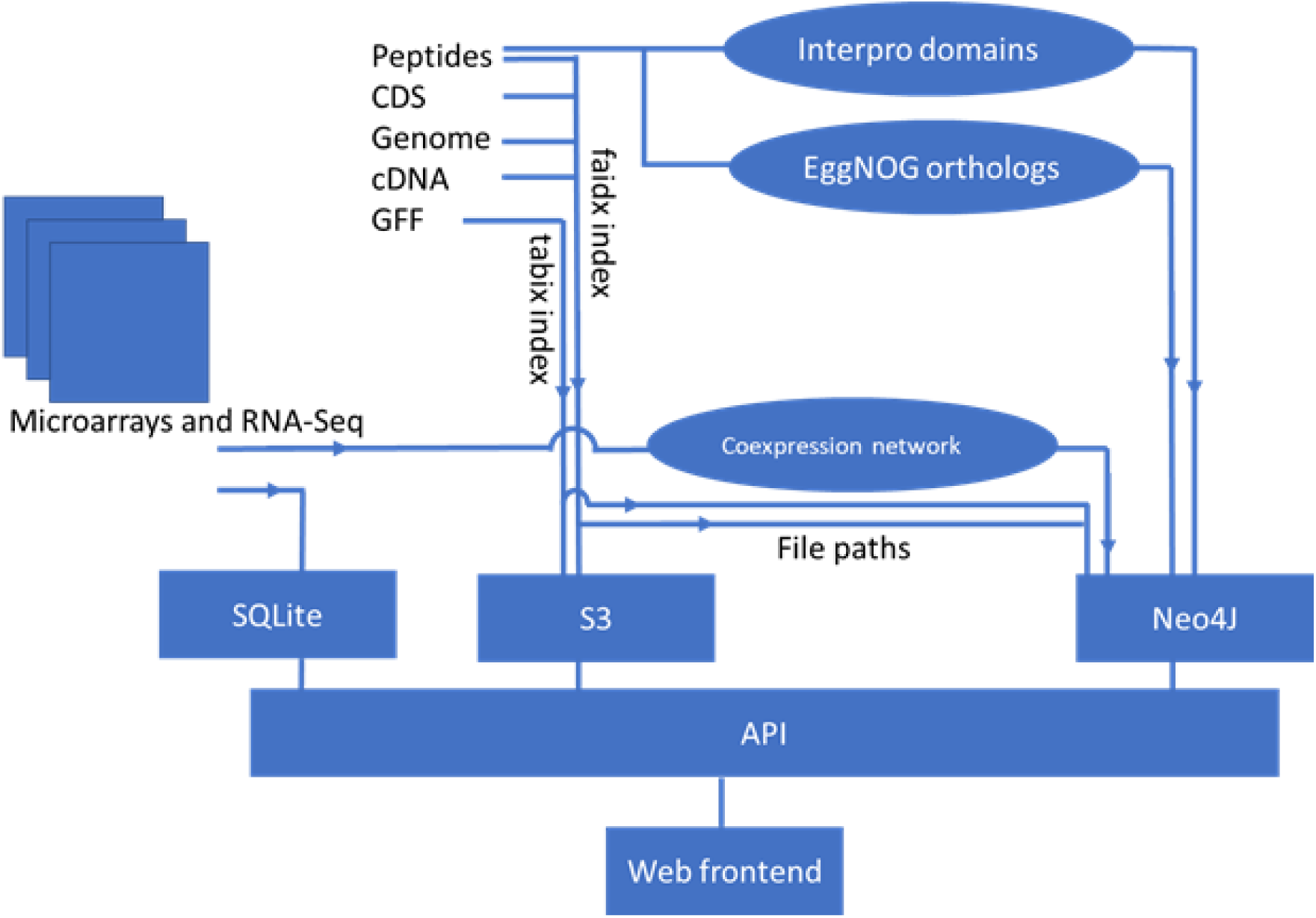
Storage and pipelines included in the data processing and storage. Microarray and RNA-seq data are stored processed and normalized (not shown). The TPMs are then stored in an SQLite database. The TPMs is also processed to generate a co-expression network, stored in Neo4J. Peptide, CDS, cDNA, Genomes and GFF files are stored in an S3 compatible object storage, together with their respective faidx and tabix indexes (Li et al., 2009; Li, 2011). Peptide sequences are processed by eggNOG (Huerta-Cepas et al., 2016) to identify ortholog families, and by InterProScan to find conserved domains. Both orthologs and domains are stored in Neo4J, connected to their respective peptides.

FASTA and GFF files, are stored using an S3 compatible object storage (Palankar et al., 2008), together with their respective indexes. Accessing this information is easy since we can traverse the graph from a given Accession to the S3 data, and use the Accession together with the file index to drag out only what we need from the flat-file, while benefiting from the features of S3 buckets, especially scalability and accessibility across servers.

To ease access to our data, we developed a web-based interface for OmicsDB::Pathogens (http://pathogens.omicsdb.org). An overview of the website is shown in Fig 3. The site is centered around gene information pages, where each page tries to summarize the knowledge about a given gene, Users can search for the genes of interest using keywords, gene identifiers or by performing a BLAST search against their sequence of interest.

**Fig 3.**
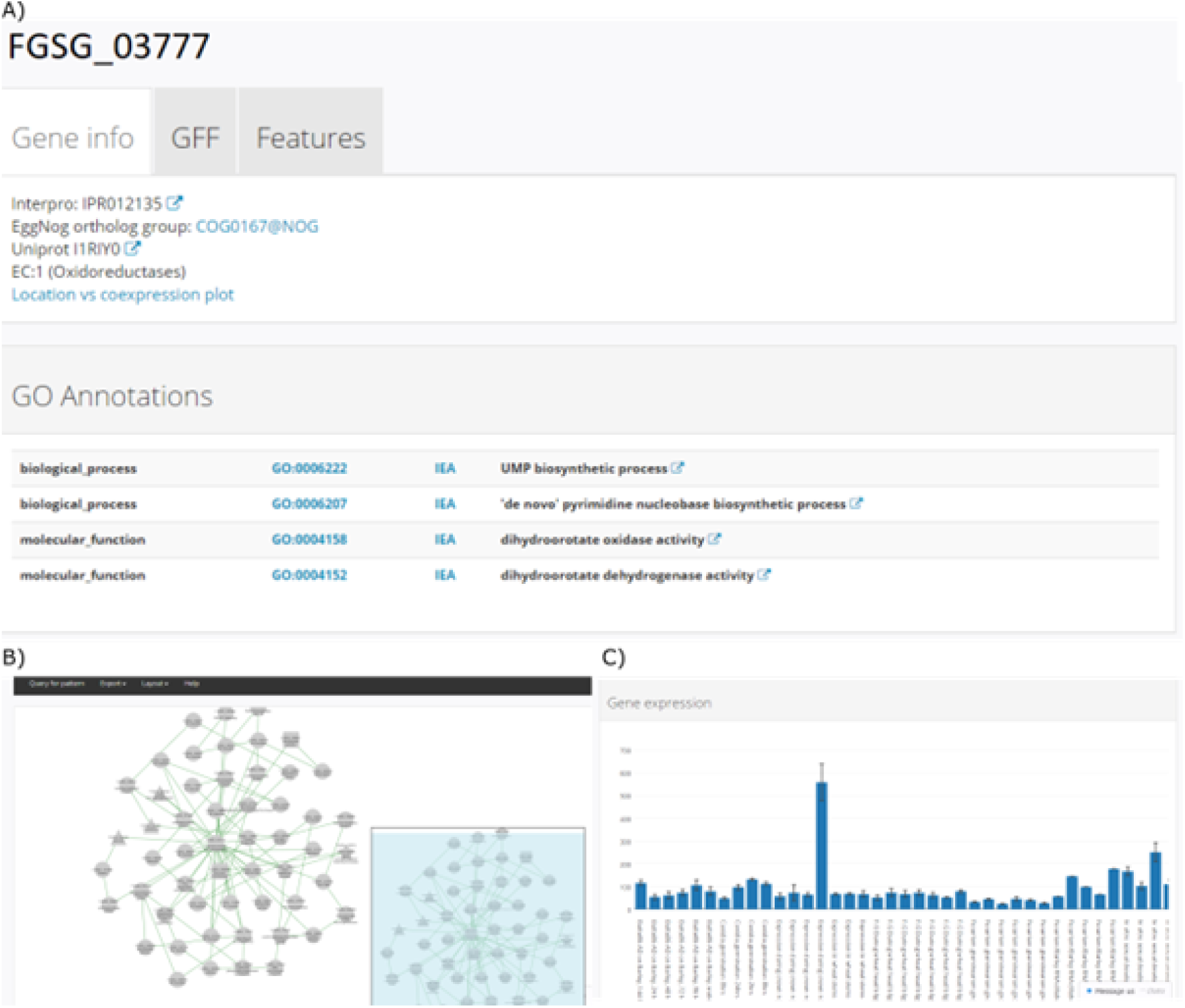
An example of the gene profile page. A) Shows the structural and functional information on the gene as well as ortholog information, possible pathways, and InterPro domains. B) Shows an example of a co-expression network. C) Shows the expression profile of a given gene. Both A, B and C are found on the same page in the browser.

### Basic functions of OmicsDB::Pathogens

The main source of information is the gene page that offers important information regarding each gene. It provides different information for each gene. An overview can be seen in Fig 3.

The top of the page (Fig 3A) shows the basic information: The gene model with intron and exon information. Gene Ontology annotations (Ashburner et al., 2000). Ortholog groups from EggNOG (Huerta-Cepas et al., 2016) and protein domains from InterProScan (Hunter et al., 2009; Jones et al., 2014) and the coding DNA sequence and the protein sequence. Further down the page (Fig 3B) is a list of co-expressed genes and a visualization of the co-expression network. As well as A bar chart of gene expression displaying the average of replicates for a given experiment (Fig 3C).

This page also serves as a gateway to explore the information within and across species. For the species without networks, this data is limited to protein domains, ortholog families, DNA and protein sequences.

The network representation of gene interactions allows the users to query for the pattern of co-regulated orthologs across other species, to see if this regulated “module” is preserved. This function can ease the annotation transfer of “biological process” terms, relying not only on sequence similarity but also on the biological context.

To give a quick overview of what types of proteins the genes in the network code for, shapes of the nodes are used. The default shape is round, but it will be given a different shape in the case of transcription factors, or if a protein is classified with one of the 1-6 Enzyme commission classifications (Cornish-Bowden, 2014). The node representing the bait gene is displayed in red. An overview of the shapes can be found in Supplementary Table 1. To represent different types of relationships, the edges are colored using the following scheme, Green edges: co-expression, red edges: physical interaction, blue edges: genetic interaction. The physical and genetic interactions are from The BioGrid database (Oughtred et al., 2019), the co-expression data is calculated using the Mutual Rank (MR) method (Obayashi et al., 2009) modified to 1/MR with a MR threshold of 50.

From the gene page is it possible to access other pages with information. Ortholog pages: Each EggNOG ortholog family has a dedicated page, with aggregated information on this family. Including genes and species in the family, functional annotations as well as papers citing the family. This enables users to easily check at a glance what information is available from other species. As well as how diverse the family is by visualizing the distribution of members per species in a pie chart. An example of an ortholog page can be seen in Fig 4. Domain pages are similar to ortholog pages, but centred around a domain, for example, PF00331 or PS51760.

**Fig 4.**
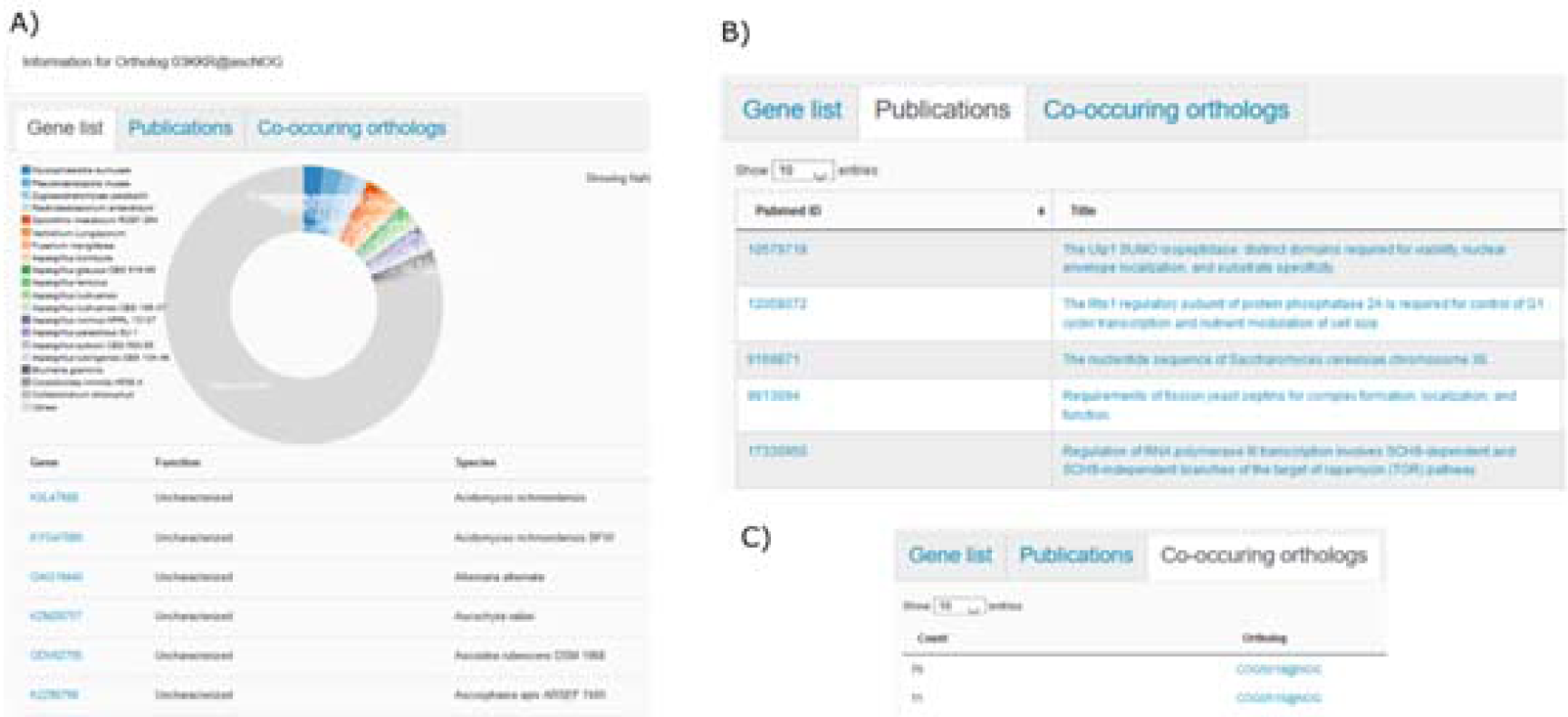
Each Ortholog has a unique page with multiple tabs. A) it shows all that has been assigned and the species they belong to. It is possible to sub select species using the pie chart. The other tabs include B) publications citing genes within this ortholog group. As well as C) orthologs that co-occur across the networks. Orthologs that often co-occur hints that they might participate in a similar process.

The Gene Ontology(Ashburner et al., 2000) has also been integrated, with pages for each term, displaying all genes annotated with a given ontology term, as well as the position of the term in the GO graph. It is thus possible to find networks in one species that resembles those affiliated with a certain ontology in another species. Traditionally the user would have had to download and annotate both datasets. However here it can be done with the click of a button.

### Advanced functions

OmicsDB also provides some tools enabling users to perform more advanced analyses. One example is cross-species network alignment. Cross-species comparisons of biological networks with interactions are still an emerging field, It allows for finding network patterns that between orthologous genes across species, hinting towards similar function. This has been used successfully in plants (Usadel et al., 2009; Movahedi et al., 2012; Hansen et al., 2014). Most work on gene function prediction using gene co-expression so far has been working on single or few species. Identifying conserved co-expression patterns across orthologs in different species can identify highly relevant candidate genes sharing similar functions or participate in the same pathway (Movahedi et al., 2012; Hansen et al., 2014).

An example of such a strategy with experimental confirmation is the study by Itkin et al., where comparative co-expression for tomato and potato was utilized, leading to the discovery of a gene cluster that is related to the steroidal glycoalkaloids pathway (Itkin et al., 2013).

It is also possible to use the surrounding network and look for enriched GO terms, using a hypergeometric test. This is done at the bulk analysis page. Where a list of genes can be analysed. A plot of the expression values for all genes in the list will be generated as well as the GO enrichment. It is also possible to export both GO enrichments as well as peptide and CDS sequences in fasta format.

### Use case: Vacuolar iron uptake

Yeast Fet3/Ftr1 (YMR056W/YER145C) are genes for two proteins that are in yeast high-affinity iron uptake at the plasma membrane (Askwith et al., 1994, 3; Stearman et al., 1996). The proteins physically connect to each other. Ftr1 is a transporter and Fet3 is an oxidoreductase. A similar system Fet5/Fth1 (YFL071W/YBR207W) exists at the vacuolar membrane and is involved in regulating vacuolar iron storage (Urbanowski and Piper, 1999)

In *Fusarium graminearum* FGSG_02143 is known to be the plasma membrane Ftr1-like protein and the homologous protein FGSG_05160 is probably the Fth1-like protein located at the vacuole membrane. To test if this can be the case FGSG_05160 was used as bait and the result is displayed together with the networks for yeast Ftr1 and Fth1. The aligned network can be seen in Fig. 5. Red edges represent possible orthologs and black edges co-expression within the species. The subcellular location for some genes products in Yeast has been marked.

**Fig 5.**
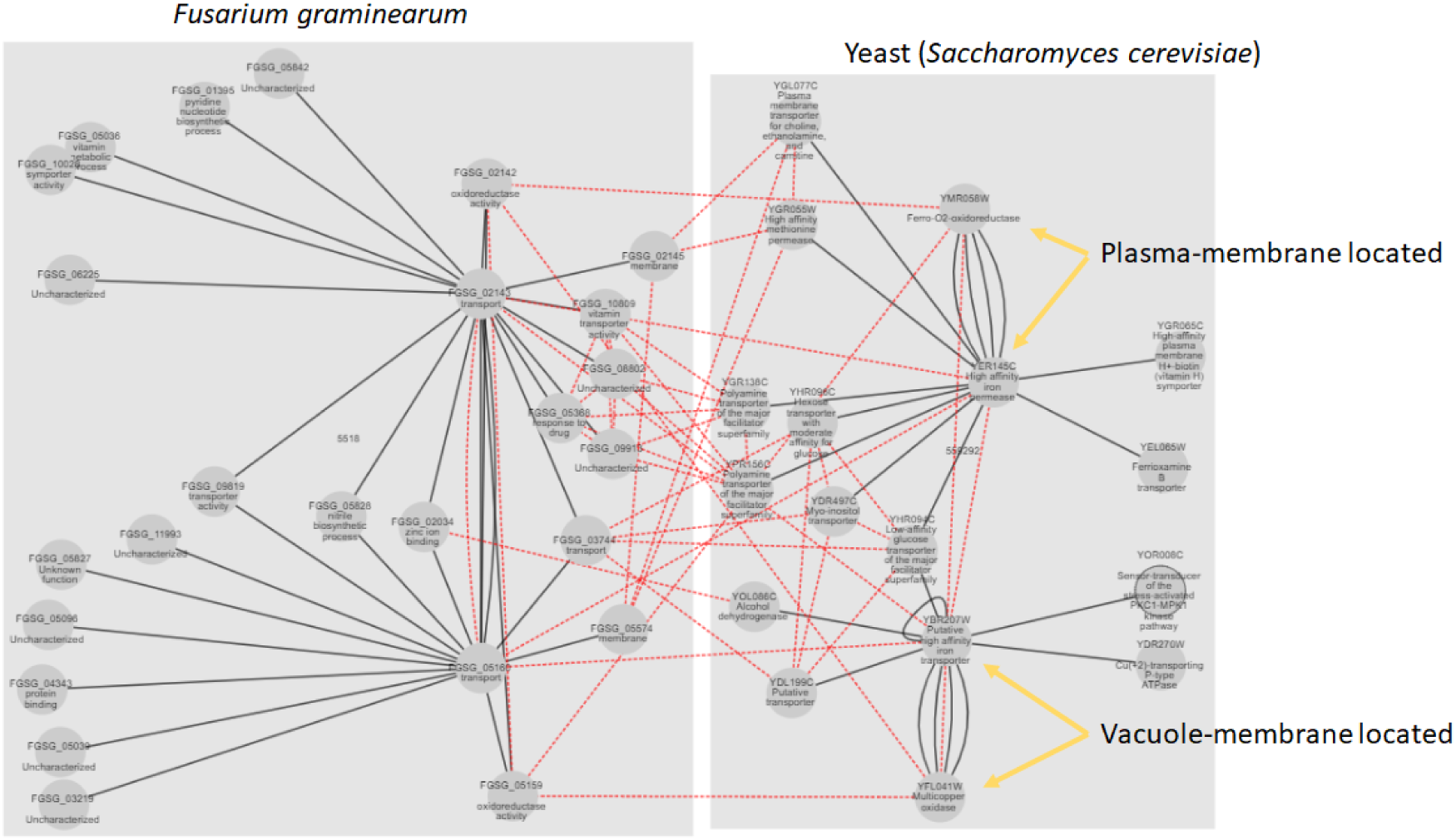
The alignment of the co-expression neighbourhoods from Yeast and F. graminearum, Using FGSG_05160 as a bait, and searching for counterparts in Yeast with similar co-expression patterns. Black edges represent co-expression edges, red edges represent possible orthologs relationship. It can be seen that large parts of the network are mirrored across the organisms, hinting towards the genes carrying out the same functionality.

What can be seen is that the vacuolar and the plasma-membrane iron transport is more co-regulated in the *F. graminearum* data than in the yeast data (direct strong co-regulation between FGSG_05160 and FGSG_02143). The pattern of co-regulation of the other proteins with orthologues in yeast also supports that FGSG_05160 is a vacuolar located Fth1-orthologue. This has, however, to be confirmed by direct experiments.

## Conclusions and future directions

In this study, we developed a system for handling biological omics data. We processed gene expression data, as well as generated co-expression networks for 10 species.

We established an interactive web interface omicsdb.org to provide access to the data as well as the analysis platforms for the public. We plan to continue to improve the quality and functionality of this database, by regularly updating with new publicly available data.

Currently the database contains networks for 4 pathogens and 6 references, however, there is a growing need for understanding how plant pathogens work, to alleviate this need we will include new species as when they become available.

## Materials and methods

### External tools

BLAST+ (Camacho et al., 2009) has been integrated to enable the users to easily find their gene of interest, allowing the user to BLAST their bait sequence against our database. This is practical in case they have identifiers not present in our database, or the user working on species not yet included.

### Construction of gene co-expression data

For the construction of co-expression database for plant pathogens, we selected 4 pathogenic species (*Fusarium graminearum, Ustilago maydis, Blumeria graminis, Neurospora crassa*), based on the availability of gene expression data, as well as 6 reference species (*Escherichia coli, Schizosaccharomyces pombe, Saccharomyces cerevisiae, Arabidopsis thaliana, Mus musculus, Homo sapiens*).

Expression data were downloaded from Arrayexpress (Kolesnikov et al., 2015), and normalized using the following methods: For Agilent data, the R-package limma (Ritchie et al., 2015) were used. For Affymetrix, Affymetrix Powertools v. was used, running the RMA algorithm. For NimbleGen, DEVA V 1.02 (Roche) was used. RNA-Seq data were subjected to QC using FastQC (Andrews), reads were mapped using TopHat2 (Kim et al., 2013), counted using HTseq-count (Anders et al., 2015) and normalized using the VST algorithm implemented in DESeq2 (Love et al., 2014), run in R v. 3.4 (R Core Team, 2013).

Genome annotations were derived from Pombase for *S. pombe* (Wood et al., 2012), SGD for *S.* c*erevisiae* (Cherry et al., 1998), and Ensembl for the remaining species. Transcription factor annotations were downloaded from (Liu et al., 2013), Enzyme Commision numbers were obtained from Uniprot (The Uniprot Consortium, 2019). Biological, chemical and genetic interactions were derived from BioGrid (Oughtred et al., 2019).

The Mutual Rank value of the weighted Pearson’s correlation coefficient was used as the measure of co-expression, as described by (Obayashi et al., 2009). A threshold of 50 was applied. To make it comparable with protein interactions, the MR was normalized by taking 1/MR ensuring that 1 was equivalent to the best value. Mutual Rank is calculated as

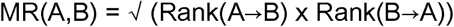

With A and B represent genes. GFF files were stored in tabix format (Li, 2011) peptide and DNA sequences were stored in faidx format implemented in Samtools (Li et al., 2009).

### Functional Annotations

Protein domains including PFAM domains (Finn et al., 2014) and Panther (Mi et al., 2013) were identified using InterProScan v. 5.29-68.0 (Hunter et al., 2009; Jones et al., 2014),

Ortholog groups were identified using EggNOG (Huerta-Cepas et al., 2016), which also provided predicted GO terms. For *E. coli, S. pombe, S. cerevisiae A. thaliana, M. musculus and Homo Sapiens*, experimentally validated GO terms could be downloaded from the GeneOntology website (The Gene Ontology Consortium, 2019). The Gene Ontology website also provides mappings from domains identified by InterProScan to their associated GO terms. These maps were used to further improve the annotations.

Papers citing a given gene or gene family were retrieved from Uniprot (The Uniprot Consortium, 2019).

For reference, Arabidopsis thaliana, Mus musculus and Homo sapiens were included, annotations were derived similarly to the pathogens.

### Network alignment

The network alignment uses the “@NOG” ortholog families, if you provide a bait gene, it will query all other genes from this family, and compare how many orthologs are shared in the 1st-degree neighbourhood. To calculate how much of the network around two genes is similar, the Jaccard index for the gene families shared across the networks is then calculated following

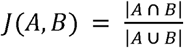

Where A and B represent the gene families in the network surrounding your bait gene and the gene it is compared against respectively. The numerator is the intersection of gene families, and denominator represents the union of all gene families in the 1st-degree neighbourhood.

### Ontology Enrichment

Enrichment of Gene Ontology terms in sets of genes is calculated using the Hypergeometric distribution function, implemented in Scipy (Virtanen et al., 2020). For the calculation of background terms, following the true path rule, all ancestors of any given annotation are considered.

### Database implementation

The system runs on Linux v. 18.04 LTS Bionic Beaver. The web-service was implemented in a Python-based web application framework, Flask v. 1.0.2, with SQLite and Neo4J 3.4 (www.neo4j.com) as the backend databases for the expression data and everything else respectively. The website is being served using gunicorn and NGINX. The co-expressed gene networks as well as the directed acyclic graphs for the GO terms were visualized using Cytoscape.js v. 3.14 (Franz et al., 2016), gene expression plots were generated using dc.js (https://dc-js.github.io/dc.js/). For the HTLM page layout, the bootstrap framework was used (www.getbootstrap.com), and for general purpose usability features the Jquery.js library (https://jquery.com) was used

BLAST+ (Camacho et al., 2009) was installed and a custom HTML interface was developed. Icons were obtained from Font Awesome v. 5 (www.fontawesome.com).

## Author Contributions

BOH and SO designed the study, BOH developed the database and processed the data. BOH and SO wrote the manuscript

