## Supplementary data for "OmicsDB::Pathogens - A database for exploring functional networks of plant pathogens" (from another version of this preprint: version 1, 2020-03-19)

### Table 1. Species with networks

| ***Species*** | **Coding sequence source** | **No genes covered in database** | **Number of edges** | **Number of samples** |
| --- | --- | --- | --- | --- |
| *Fusarium graminearum* | (King et al., 2015) | 13317 | 429207 | 202 |
| *Ustilago maydis* | (Kämper et al., 2006) | 6781 | 232268 | 236 |
| *Blumeria graminis* | (Wicker et al., 2013) | 6470 | 33960 | 68 |
| *Neurospora crassa* | (Galagan et al., 2003) | 9730 | 131132 | 102 |
| *Schizosaccharomyces pombe* | (Wood et al., 2002) | 5134 | 250865 | 265 |
| *Saccharomyces cerevisiae* | (Giaever et al., 2002) | 5887 | 1533434 | 1123 |
| *Escherichia coli* | (The Complete Genome Sequence of Escherichia coli K-12 \| Science) | 3212 | 383284 | 701 |

The source of the genome, the number of gene that are covered in our database, the number of co-expression edges and the number of transcriptomics samples used for generating the network

### Table 2. Legend to network

| **Shape** |  | **Description** |
| --- | --- | --- |
| Star | 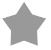 | Transcription factor |
| Octagons | 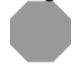 | EC:1 (Oxidoreductases): |
| Round rectangle | 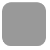 | EC:2 (Transferases): |
| Triangle | 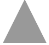 | EC:3 (Hydrolases): |
| Rectangle | 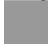 | EC:4 (Lyases): |
| Pentagon | 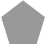 | EC:5 (Isomerases): |
| Heptagon | 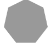 | EC:6 (Ligases) |
| Round | 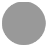 | Others or genes without info or not in the classes above |
